# Phase-free local ancestry inference mitigates the impact of switch errors on phase-based methods

**DOI:** 10.1101/2023.12.02.569669

**Authors:** Siddharth Avadhanam, Amy L. Williams

## Abstract

Local ancestry inference (LAI) is an indispensable component of a variety of analyses in medical and population genetics, from admixture mapping to characterizing demographic history. However, the accuracy of LAI depends on a number of factors such as phase quality (for phase-based LAI methods), time since admixture of the population under study, and other factors. Here we present an empirical analysis of four LAI methods using simulated individuals of mixed African and European ancestry, examining the impact of variable phase quality and a range of demographic scenarios. We found that regardless of phasing options, calls from LAI methods that operate on unphased genotypes (phase-free LAI) have 2.6-4.6% higher Pearson correlation with the ground truth than methods that operate on phased genotypes (phase-based LAI). Applying the TRACTOR phase-correction algorithm led to modest improvements in phase-based LAI, but despite this, the Pearson correlation of phase-free LAI remained 2.4-3.8% higher than phase-corrected phase-based approaches (considering the best performing methods in each category). Phase-free and phase-based LAI accuracy differences can dramatically impact downstream analyses: estimates of the time since admixture using phase-based LAI tracts are upwardly biased by ≈10 generations using our highest quality phased data but have virtually no bias using phase-free LAI calls. Our study underscores the strong dependence of phase-based LAI accuracy on phase quality and highlights the merits of LAI approaches that analyze unphased genetic data.

## Introduction

The problem of inferring the ancestral population of each locus in an admixed individual’s genome, or local ancestry inference (LAI), has received attention for well over a decade ^1–9^, and a variety of LAI approaches now exist. Early LAI methods only analyzed markers that are not in linkage disequilibrium (LD) ^1–3^, thus reducing the information available for inference while greatly simplifying the modeling. The advent of the Li and Stephens haplotype model ^10^, which accounts for LD, enabled the development of several LAI methods that leverage all available marker data. HAPMIX ^5^ was one of the first approaches to leverage this rich information, which yielded dramatic improvements in both the accuracy and resolution of LAI compared to earlier methods. Subsequent methods that allow for multiple source populations or have improved runtime scaling have been developed ^6,7,9^, with many recent LAI methods requiring that the admixed individuals be phased prior to analysis ^7,9^, while HAPMIX, LAMP-LD ^6^, and ELAI ^8^ analyze unphased genotypes and output inferred local ancestry in an unphased manner.

Local ancestry calls have wide-ranging use in both medical and population genetics. Admixture mapping ^11,12^ detects trait-associated loci by exploiting the fact that inferred local ancestry tracts are a proxy for variants not directly tested in a study and for a large number of haplotypic backgrounds from the same population. Moreover, methods exist for increasing the power of genome-wide association studies ^13,14^ and for improving heritability estimates and mapping in eQTL studies ^15^ by including local ancestry calls. At the same time, population genetic analyses have used LAI calls to characterize demographic patterns among admixed groups, detecting geographic trends that reflect historical migrations ^16^, and have inferred times since admixture using local ancestry tract lengths ^17^.

Phasing errors, or mis-assignment of alleles to haplotypes (herein measured as switch errors) can confound LAI by introducing a short tract of a different ancestry onto a haplotype (in cases of two nearby switch errors) or by prematurely ending a tract. Besides merely switching the ancestry assignment to the opposite haplotype, switch errors decrease the quality of LAI calls in part because accurately detecting tract boundaries is difficult, and short, incorrectly phased tracts may not be detected. Furthermore, LAI switch errors can have important consequences on inferring admixture demography in that they alter observed tract lengths ^17^. Phase-based LAI methods sometimes attempt to account for phasing errors, as in RFMix, which jointly models switch errors and local ancestry ^7^. By contrast, phase-free LAI methods do not read phase information for the target samples and are therefore unaffected by switch errors in these individuals.

Given the many applications of LAI and an appreciation of the associated phasing issues, characterizing the performance of LAI methods is important; yet many comparisons are in the papers that describe new methods, with relatively few independent analyses. One recent LAI benchmarking study included a range of LAI methods and is complementary to our work in that it does not analyze the impact of phasing errors on the tools ^18^.

Here we present a comparison of four LAI methods using genome-wide data for simulated individuals of mixed African and European ancestry. To appropriately model phasing error, we converted the simulated haplotypes into diploid genotypes and phased these using a range of sample sizes, thus accounting for and quantifying the impact of phasing errors on LAI. Moreover, we considered the impact of demography— genome-wide ancestry fraction and time since admixture—as well as phasing choice with respect to the LAI reference panels on LAI accuracy.

## Methods

To systematically investigate the impact of various demographic parameters and phasing options on LAI accuracy, we developed a pipeline (Figure 1) that: 1) simulates genotypes with mixed African and European ancestry for African ancestry proportion *p* ∈ {0.2, 0.5, 0.8} and time since admixture *T* ∈ {5, 6, 7}; and 2) phases these genotypes using three different sample sizes, which substantially impacts phase quality. At the same time, we varied phasing with respect to the unadmixed reference panels by three different panel phasing strategies (see below).

**Figure 1:**
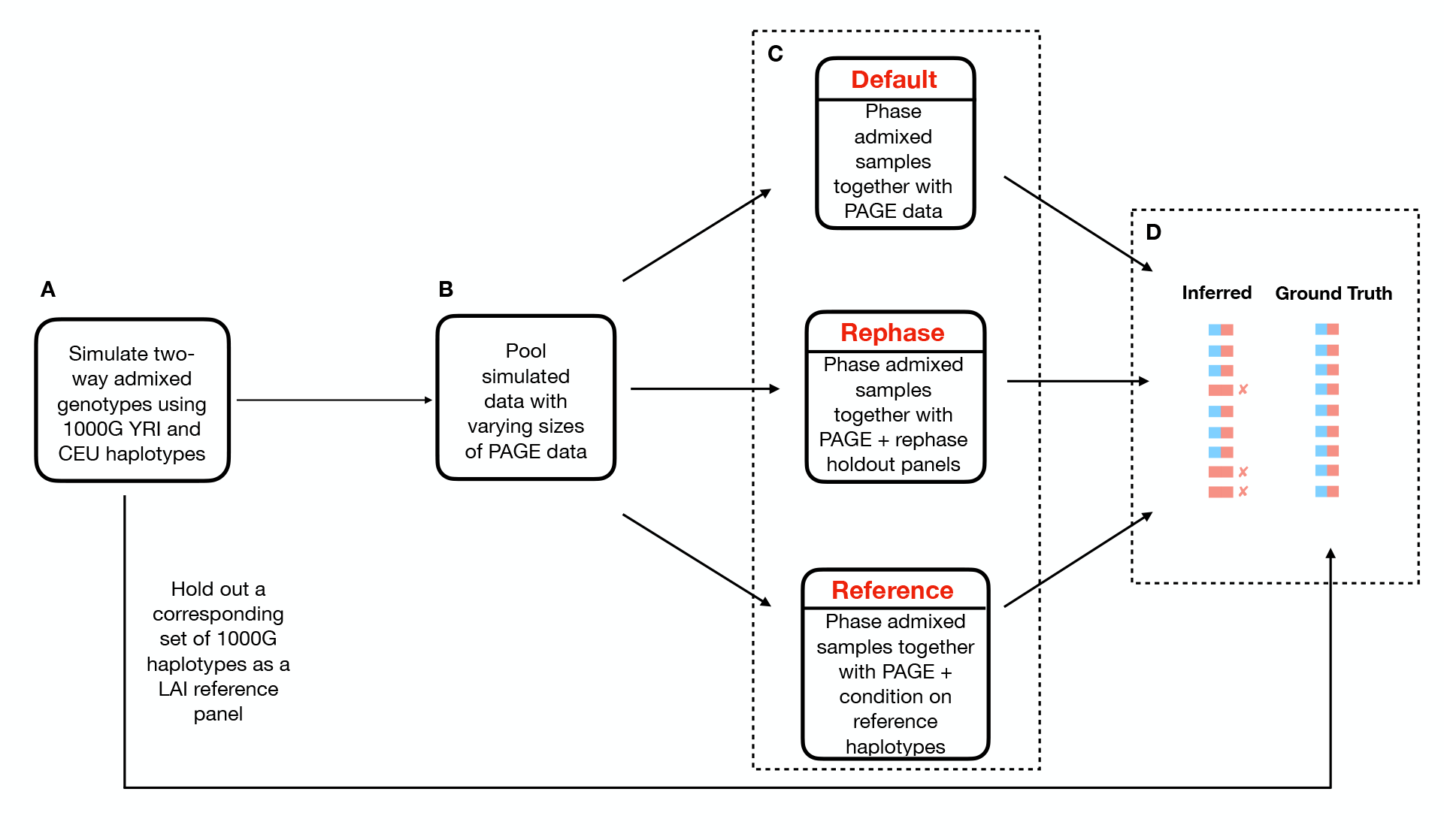
Flowchart depicting our simulation and phasing pipeline. (A) First, we simulated two-way admixed individuals under different settings of *T* and *p*, and we held out some unadmixed individuals from the simulation founders for use as LAI reference panels. (B) Next we pooled the simulated data from (A) with small, medium, and large sample sizes from the PAGE data. (C) Third, we applied three different panel phasing strategies termed default, rephase, and reference. (D) Finally, we compared the true and inferred local ancestry calls using Pearson correlation.

The LAI methods we benchmarked fall in two categories: phase-free, which take unphased data as input and output unphased local ancestry calls, and phase-based, which require phased haplotypes and output phased local ancestry calls. In the phase-free category, we applied HAPMIX (version 2) and LAMP-LD (version 1.1), and in the phase-based category we applied RFMix (version 2), and FLARE (version 0.3.0).

The different phasing options we applied reflect workflows that could occur under variable real-world constraints— namely access to admixed datasets of different sizes, choice of panel-based phasing strategy, and choice of LAI method (which may itself be constrained by phase quality). For example, a user that has few admixed samples from a population of interest might phase them jointly and retain the phase supplied by (for example) the 1000 Genomes Project (1000G) ^19^ for the unadmixed reference haplotypes (i.e., rather than rephasing this panel). Alternatively, the phasing step may not be necessary if a user applies a phase-free LAI method such as HAPMIX or LAMP-LD.

We evaluated the LAI accuracies by measuring the Pearson correlation (*R*) between the inferred and simulated local ancestry calls (the latter are the ground truth assignments) across all markers. We calculated this correlation using unphased (i.e., diploid) local ancestry calls for a two reasons. First, measuring accuracy in a phase-based way is complicated by the presence of switch errors that are introduced upstream of the LAI method. Second, ignoring phase ensures a uniform metric of comparison between phase-based and phase-free LAI methods.

Together with these accuracy calculations, we evaluated the LAI methods’ performance for estimating time since admixture using a Markovian model described by Gravel ^17^. We applied this model directly on the output of the phase-based method FLARE and indirectly using PAPI ^20^ on the output of the phase-free method HAPMIX.

### Data-efficient simulation of admixed genotypes

In recent work ^20^, we described a pipeline for simulating admixed individuals that first uses Ped-sim ^21^ to generate crossovers in a fixed pedigree, and then uses admix-simu ^22^ to sample haplotype segments at each crossover breakpoint. The latter draws each segment from the same population as the unadmixed ancestor that transmitted it, with the advantage (versus simply using Ped-sim) of requiring far fewer unadmixed individuals to simulate each person. We employed a pedigree topology containing the admixed individual in the most recent generation (generation 0) and all that person’s ancestors up to generation *T* included (i.e., with 2^*T*^ founders in the oldest generation). The pipeline also uses *p* to specify the proportion of African founders.

For each of the nine combined settings of *T* and *p*, we simulated a batch of 30 admixed individuals using 35 YRI and 31 CEU founders. These founders are drawn from a set of 107 YRI and 95 CEU 1000G genotypes (where we excluded all trio/duo children). We chose these values so as to maximize the number of founders used to simulate each batch while further ensuring that a larger non-overlapping hold out set of 72 YRI and 64 CEU individuals were available for use as LAI reference panels (for each simulation batch). We ran the same simulation in triplicate, where each replicate operates on the same set of founders but re-samples crossover breakpoints and the ancestral segments copied from. This gives us a total of 90 admixed individuals for each combined setting of *T* and *p*.

For each batch (including replicates) we merged the 30 simulated samples with additional admixed individuals of three different phasing sample sizes and applied three different panel phasing strategies (see the next subsection and Figure 1) yielding a total of nine phasing conditions analyzed for each combined setting of *p* and *T*. A benefit of this is that every comparison between phasing sample size and panel strategy operate on the same set of data, thus automatically controlling for the noise generated by the simulation of admixed genotypes. In other words, while comparisons between different values of *p* and/or *T* analyze different simulated admixed samples, comparisons between different phasing sample sizes or panel phasing strategies (but the same *p* and *T*) consider the exact same simulated data.

### Varying simulation phase quality and panel phasing strategies

We induced varying degrees of phase quality by pooling the simulated admixed individuals with variable numbers of genotypes from the BioMe Biobank subset of the Population Architecture using Genomics and Epidemiology (PAGE II; dbGaP:phs000925.v1.p1)^23^ study and phasing them jointly using SHAPEIT4^24^ (see below and Figure 1). To that end, we ascertained two-way admixed PAGE individuals of mixed African and European ancestry by following a procedure that we employed in earlier work ^20^. We first ran ADMIXTURE ^25^ with K=5 on the PAGE samples merged with 176 HapMap trio parents evenly split between the CEU and YRI populations; this allowed us to determine which of the K components corresponded to African and European ancestry. We then selected individuals with (1) ≥5% African ancestry; (2) ≥5% European ancestry; and (3) the sum of these two ancestries being ≥99.5%; this yielded 5,786 PAGE samples. We then randomly sampled 5,780 of these PAGE individuals to use as the ‘large’ sample size, 2,890 for the ‘medium’ sample size, and 580 for the ‘small’ sample size. We verified that the phasing error generally correlated inversely with sample size by measuring switch error rates using `vcftools` with the `--diff-switch-error` flag (see Results).

Besides varying the sample size, we applied three different panel-based phasing strategies: (1) the ‘default’ phasing strategy, where we retained the original 1000G phase for the panels; (2) the ‘reference’ phasing strategy, where we use the `--reference` option so that SHAPEIT4 conditions on the panel haplotypes when phasing the admixed samples (this is recommended when using such a panel for downstream analyses such as imputation or LAI); and (3) the ‘rephase’ strategy, where we pooled the reference panels with the admixed genotypes and retrieved the resulting rephased haplotypes for use as panels for LAI. The latter two strategies help make the admixed individuals’ phase more consistent with those of the reference panel, which contrasts with the default strategy, where the reference haplotypes have the phase present in the 1000G data—which were generated independent of the admixed individuals.

### Filtering and processing the PAGE data

The data we used for this work is a merging of the PAGE and 1000G data that we used previously. Full details of the merging process and quality control filters we applied are available in our earlier work ^20^, and we provide a brief summary here. The key steps of the pipeline are filtering SNPs in the PAGE dataset and intersecting these filtered data with the 1000G dataset to obtain a common set of SNPs. We filtered the PAGE SNPs using the quality control report distributed with the dataset. This applies a composite filter including filters for, among others, Hardy-Weinberg equilibrium, sites with discordant calls in duplicated samples, and those with Mendelian errors. We further ensured there were no allele coding inconsistencies between the two datasets by recoding the PAGE data to the forward strand, filtering out A/T and C/G SNPs, and applying an allele frequency difference test filter. These steps yielded 494,219 markers common to both the datsets, which we used for all analyses.

### Local ancestry inference

After phasing with SHAPEIT4, we passed the datasets as input to each of the LAI methods (in the case of phase-free LAI, we erased the phase) and we used 1000G YRI and CEU haplotypes as LAI reference panels. As described above, we ensured that in all cases the LAI reference panels and the set of individuals that we used as simulation founders were disjoint.

### Measuring LAI accuracy

We compared the inferred local ancestry calls with the ground truth by representing all markers sequentially (across all 22 chromosomes) by vectors *A*_*inf*_ and *A*_*true*_ whose elements *a* ∈ {0, 1, 2} represent the number of European haplotypes at a site. The Pearson correlation coefficient (*R*) between *A*_*inf*_ and *A*_*true*_ then gives a measure of performance for each simulation setting and LAI method used. For HAPMIX, which outputs a posterior probability of each ancestry state at each marker (i.e., *P* (*a*) for *a* ∈ {0, 1, 2}), we first calculate *E*[*A*_*inf*_] before computing the Pearson correlation coefficient between *E*[*A*_*inf*_] and *A*_*true*_. The elements of *E*[*A*_*inf*_] are *E*[*a*] = Σ *P* (*a*) *· a*, which represent the expected local ancestry call for each marker.

### Comparing admixture time estimates from phased-sensitive and phase-free LAI

Our final analysis examines the performance of estimating time since admixture using local ancestry calls from phase-free and phase-based LAI. For phase-free LAI, we applied PAPI ^20^, a tool for inferring parental admixture proportions and times since admixture from unphased local ancestry calls. PAPI produces two estimates of admixture time, one for each parent of the admixed sample, which we average to obtain a single admixture time estimate 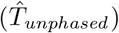.

To estimate admixture times from phased local ancestry calls, we first grouped together the tracts of Morgan length *x*_*i,a*_ corresponding to each ancestry *a* ∈ {0, 1}, where 0 ≤ *i* < *N*_*a*_, and *N*_*a*_ is the number of tracts with ancestry *a*. We then computed four statistics:

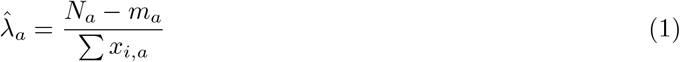

and

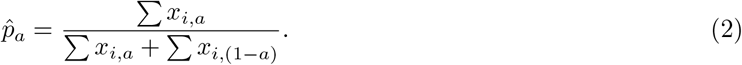

Here 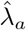 is the estimated exponential rate parameter for switching from ancestry *a* to 1 − *a*, and 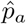 is the focal individual’s ancestry fraction from population *a*. The *m*_*a*_ term is the number of tracts of ancestry *a* that occur at the end of a chromosome (so *m*_*a*_ ≤ 22) and the −*m*_*a*_ factor in the numerator of (1) accounts for the fact that a chromosome end prematurely cuts off a tract before the next crossover is observed ^21^.

Next, we applied PAPI’s internal model for estimating admixture time to these haploid tracts. PAPI assumes that the observed local ancestry tracts are generated by transmitted crossovers within a pedigree that includes 2^*T*^ unadmixed founders of different ancestries. Further, PAPI’s model treats these founder haplotypes as a pool, where the observed haplotypes after *T* generations of crossovers are generated by a Markovian path that switches to a random haplotype in the pool at rate *T* per Morgan. Under this model, the rate of between-ancestry crossovers (or switches) from ancestry *a* to ancestry 1 − *a* is approximately the overall switch rate *T* reduced by a factor equal to the proportion of haplotypes with ancestry (1 − *a*) in the founder pool. A reasonable estimate of that proportion is simply the proportion of this ancestry in the admixed individual being analyzed, so:

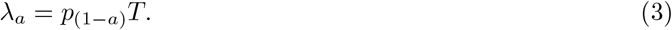

Solving this for *T* gives two estimates of admixture time, one for *a* = 0 and one for *a* = 1: 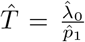 and 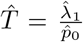. We combined these by simply averaging them, and our estimate of admixture time from phased LAI is thus 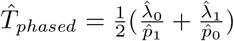.

## Results

We measured LAI accuracy for two phase-free methods—HAPMIX and LAMP-LD—and two phase-based methods—RFMix and FLARE—across every combination of four input parameters: *T* ∈ {5, 6, 7}, *p* ∈ {0.2, 0.5, 0.8}, phasing sample size (small, medium and large), and panel phasing strategy (default, reference, and rephase) (see Methods). Prior to these LAI analyses, we calculated switch error rates in the phased data in order to investigate the sensitivity of phase quality to the parameters; *a priori* we expected only phasing sample size to have a strong effect on quality. Further, we examined the potential performance of phase correction algorithms that rely on phased local ancestry calls to find and correct switch errors in the input to phase-based LAI. Specifically, we evaluated the impact of TRACTOR’s ^14^ phase correction algorithm on RFMix’s accuracy. Finally, we estimated the impact of LAI accuracy on downstream admixture time estimates using LAI tracts from one phase-free method (HAPMIX) and one phase-based method (FLARE).

### Input sample size and other simulation parameters affect switch error rates

Following phasing with SHAPEIT4, we measured switch error rates in the simulated samples for every setting of *T, p*, sample size, and panel phasing strategy (Figure 2, S4). Overall, the variable with the largest impact on phase quality is sample size. Regardless of strategy or simulation setting, jointly phasing the simulated individuals with the largest possible set of accompanying samples produced the lowest switch error rates, as is commonly seen in phasing analyses ^26^(Figure 2C). Averaged across all settings of *T, p*, and phasing method, the switch error rates for the small, medium, and large phasing sample sizes are 4.65%, 3.37%, and 3.01%, respectively. The improvement obtained by moving from the small (*N* = 580) to medium (*N* = 2, 890) sample size is considerably greater (27.5%) than that of moving from the medium to large (*N* = 5, 780) sample size (10.7%). This is again consistent with prior studies: phase quality improves dramatically when increasing from smaller sample sizes, but shows a trend of diminishing returns in larger samples ^26^.

**Figure 2:**
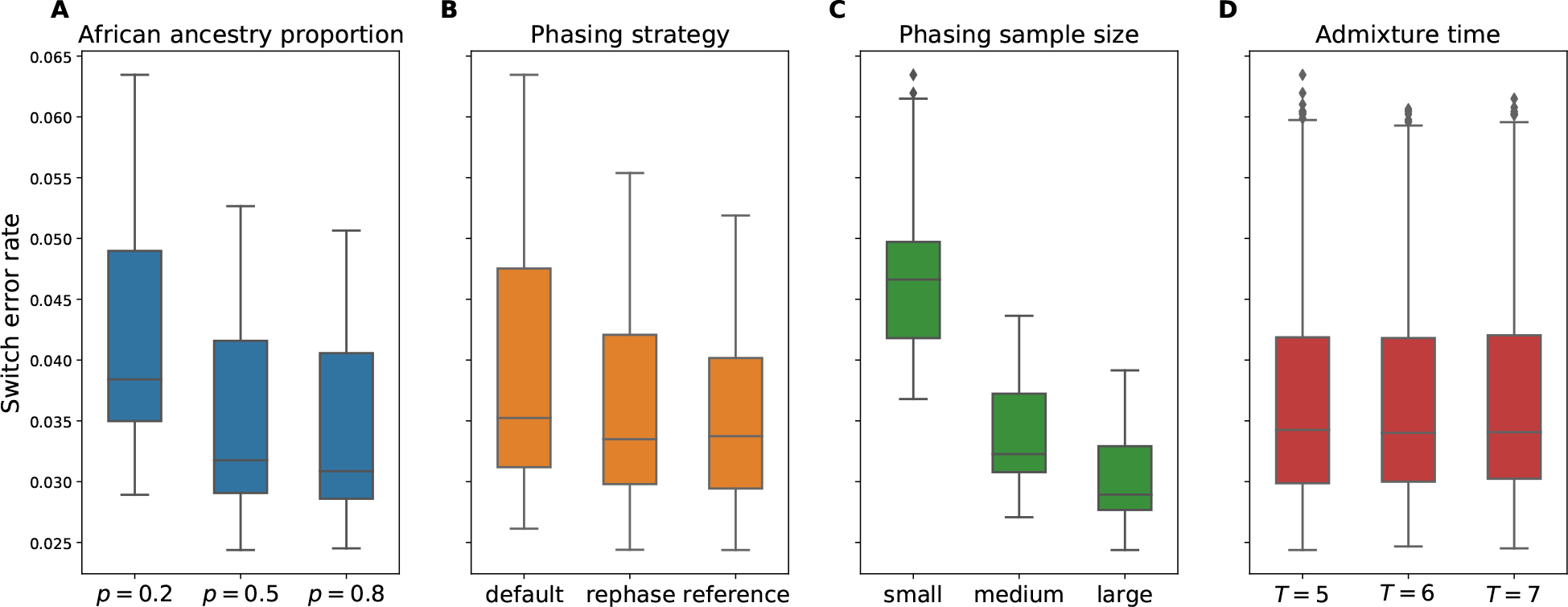
Switch error rates depend on simulation parameters. Switch error rates plotted against (A) proportion of African ancestry *p*, (B) panel phasing strategy, (C) phasing sample size, and (D) admixture time *T* (i.e., generations since admixture). Each panel includes data points from all other simulation variables.

As may be expected, phase quality is also sensitive to panel phasing strategy (see Figure 2B, S1): the average switch error rate is lowest for the reference phasing method at 3.52%, followed by the rephase method at 3.61%, and default phasing at 3.90%. This indicates that including unadmixed reference haplotypes when phasing—either by pooling them with the admixed individuals as in the rephase approach, or conditioning on them as in reference phasing—improves phase quality markedly. Our analyses, however, are consistent with the improvement in the reference and rephase strategies being at least partly due to the increase in the effective phasing sample size with the inclusion of additional haplotypes.

In turn, phasing simulated individuals with a larger proportion of African ancestry (*p*) yields improved phase, with the lowest switch error rate of 3.40% occurring when *p* = 0.8, followed by 3.48% when *p* = 0.5, and 4.15% when *p* = 0.2 (see Figure 2A, S2). A possible explanation is that admixed haplotypes simulated with *p* = 0.8 may be more similar to the admixed haplotypes from the PAGE dataset that they are pooled with: the PAGE dataset has a per-person average *p* = 0.71^20^. Finally, switch error rates show effectively no dependence on the time since admixture *T*, with only slight variation that may be driven by statistical noise (3.68%, 3.67%, and 3.69%, respectively, for *T* = 5, 6, and 7; Figure 2D).

### Phase-free LAI is more accurate than phase-based LAI

A central question in our analysis is whether phase-free LAI is more accurate than phase-based LAI given the phase quality our data provided. We found that both phase-free methods (HAPMIX and LAMP-LD) are substantially more accurate than the phase-based methods (RFMix and FLARE), even when the latter are given the highest quality phased data (Figure 3, S3). In particular, HAPMIX outperforms all the other methods across all parameter settings, with a average *R* of 0.988 (range 0.965 − 0.995), with LAMP-LD’s average *R* being slightly lower at 0.978 (range 0.946 − 0.993). Corresponding *R* values for the phase-based methods are 0.956 for FLARE (range 0.905 − 0.983) and 0.957 for RFMix (range 0.909 − 0.981).

**Figure 3:**
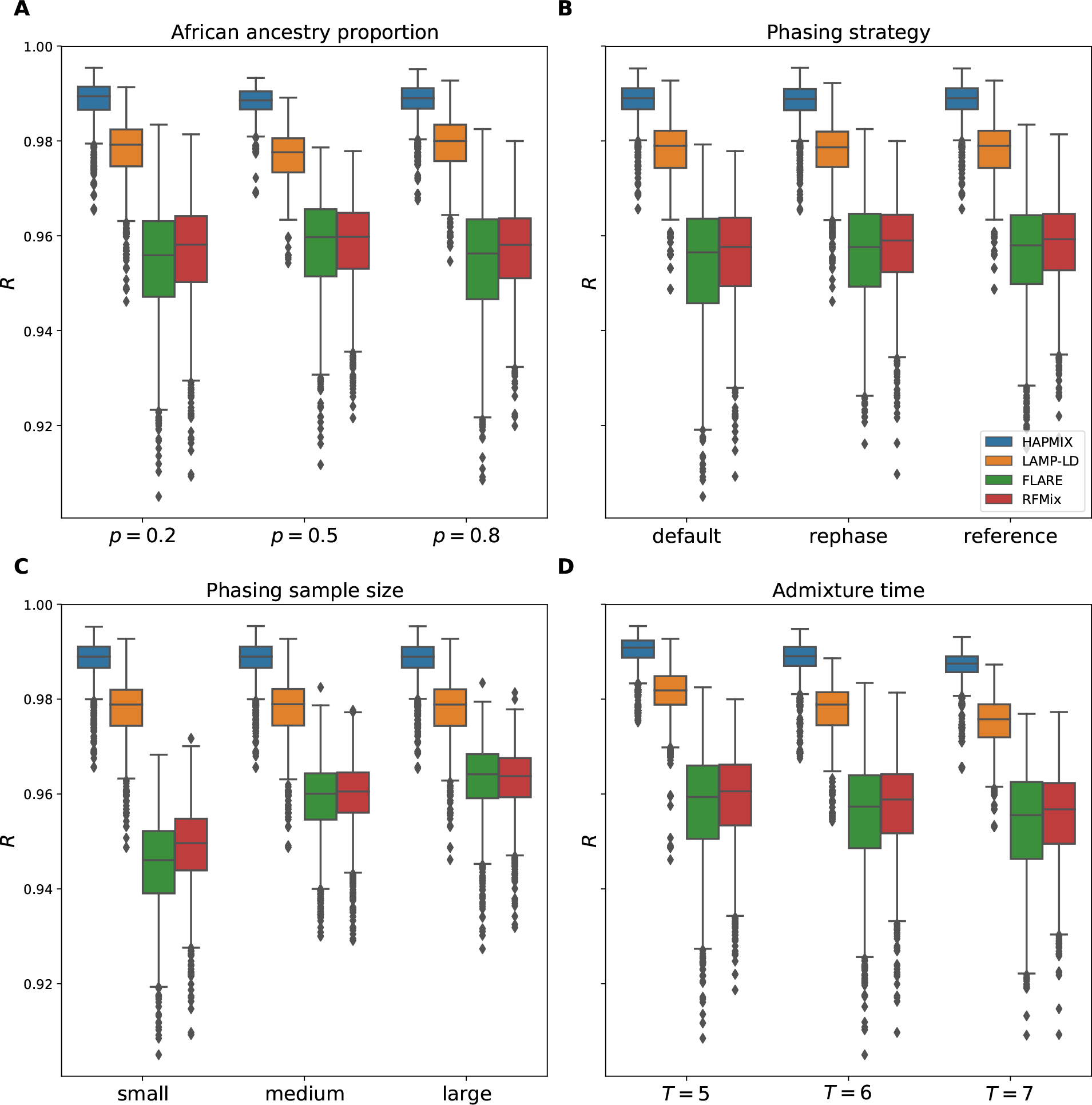
Performance of local ancestry inference methods across all simulation parameters. Correlations between inferred and true local ancestry assignments *R* for all LAI methods plotted against (A) proportion of African ancestry *p*, (B) panel phasing strategy, (C) phasing sample size, and (D) admixture time *T* (i.e., generations since admixture). Each panel includes data points from all other simulation variables.

A further advantage of using phase-free methods is that they ignore the phase of the target samples by design and are therefore robust to the impact of phasing-specific variables (notably we input the same data to all tools, but erased the phase for the phase-free methods). In particular, HAPMIX and LAMP-LD show an average *R* that is identical (*R* = 0.988 and 0.978, respectively) across all sample sizes and for the default and reference phasing strategies. The *R* values for HAPMIX and LAMP-LD are nearly identical for the rephase phasing strategy, varying at the fourth significant figure (results not shown). As mentioned previously, comparisons between sample sizes or phasing strategies are automatically noise-controlled because they consider the exact same simulated individuals (see Methods).

### Phasing strategies and phasing sample sizes strongly impact phase-based LAI performance

As would be expected for phase-based LAI approaches, RFMix and FLARE are both sensitive to phasing sample size (Figure 3C, S3), with RFMix performing identically to FLARE for medium (*R* = 0.959) and large (*R* = 0.963) sample sizes, but outperforming FLARE when the sample size is low (*R* = 0.949 and 0.945 respectively). Both methods underperform relative to the phase-free methods, as noted above.

With phase-based LAI, we find that the reference and rephase strategies work equally well and are more accurate than the default strategy (Figure 3D, S3). Specifically, RFMix has an accuracy of *R* = 0.958 for both the reference and rephase approaches (identical up to three significant figures as before) and *R* = 0.956 for the default strategy. FLARE has *R* = 0.956 for the reference and rephase strategies, and an *R* value of 0.954 with the default approach. These panel phasing strategies are fairly comparable broken down this way, and RFMix has a slight advantage over FLARE in these settings.

When the input phasing sample sizes are small, the improvements of reference and rephase strategies (*R* = 0.951 and 0.950 respectively for RFMix, and 0.947 for both methods using FLARE) over the default phasing method (*R* = 0.946 and 0.941 for RFMix and FLARE, respectively) are more substantial. RFMix has a slight advantage in terms of overall accuracy, and a more modest drop in performance with respect to sample size regardless of phasing strategy.

### HAPMIX is comparatively more robust to variation in demographic parameters

All LAI methods that we examined are relatively insensitive to African admixture proportion *p*. For instance, the phase-based methods RFMix and FLARE have virtually identical (up to three significant figures) *R* of 0.957 and 0.954, respectively, at *p* = 0.2 and *p* = 0.8, and a slightly higher *R* of 0.959 and 0.958, respectively, when *p* = 0.5 (Figure 3A, S3). LAMP-LD has a dip in accuracy between *p* = 0.2 and *p* = 0.5, from 0.978 to 0.976, and an increase in *R* to 0.979 when *p* = 0.8. Overall, HAPMIX is the most robust to changes in *p*, staying consistently more accurate than the rest at *R* = 0.988 (identically up to three significant figures) for all values of *p*.

In contrast to *p*, all methods show a similar and clear trend of decreasing accuracy as *T* increases (Figure 3D, S3). More specifically, when moving from *T* = 5 to *T* = 7, *R* decreases by 0.003 for HAPMIX, 0.006 for LAMP-LD, and by 0.004 for each of FLARE and RFMix. This is likely due to the increased number of ancestral recombinations for larger values of *T* that produce shorter local ancestry tracts. As shorter tracts have fewer SNPs than longer ones, they have lower informativeness, making LAI more challenging. Furthermore, calling local ancestry at tract boundaries is difficult, and a larger number of such boundaries may also contribute to the reduction in accuracy. Here, as with the admixture proportion, HAPMIX proves to be the most robust LAI method with respect to changes in demography.

### Phase-based LAI remains less accurate than phase-free LAI after phase correction

TRACTOR’s phase correction algorithm ^14^ rephases input data by detecting local ancestry-based switch errors, i.e., nearby local ancestry calls where the ancestries of the two haplotypes switch. We examined the correlation of RFMix’s local ancestry calls with the ground truth after applying TRACTOR, finding modest improvements of *R* = 0.960 using the corrected haplotypes versus *R* = 0.957 with the original data (averaged over all simulation parameters; see Figure 4). With the exception of the default panel phasing strategy, we see improvement across all sample sizes, demographic parameters, and phasing conditions (Figure S5). Despite these improvements, the accuracy of phase-based LAI remains much lower than that of phase-free LAI. For example, the average *R* for HAPMIX is 0.988, which is still a 2.92% relative improvement over TRACTOR-corrected RFMix (*R* = 0.960).

**Figure 4:**
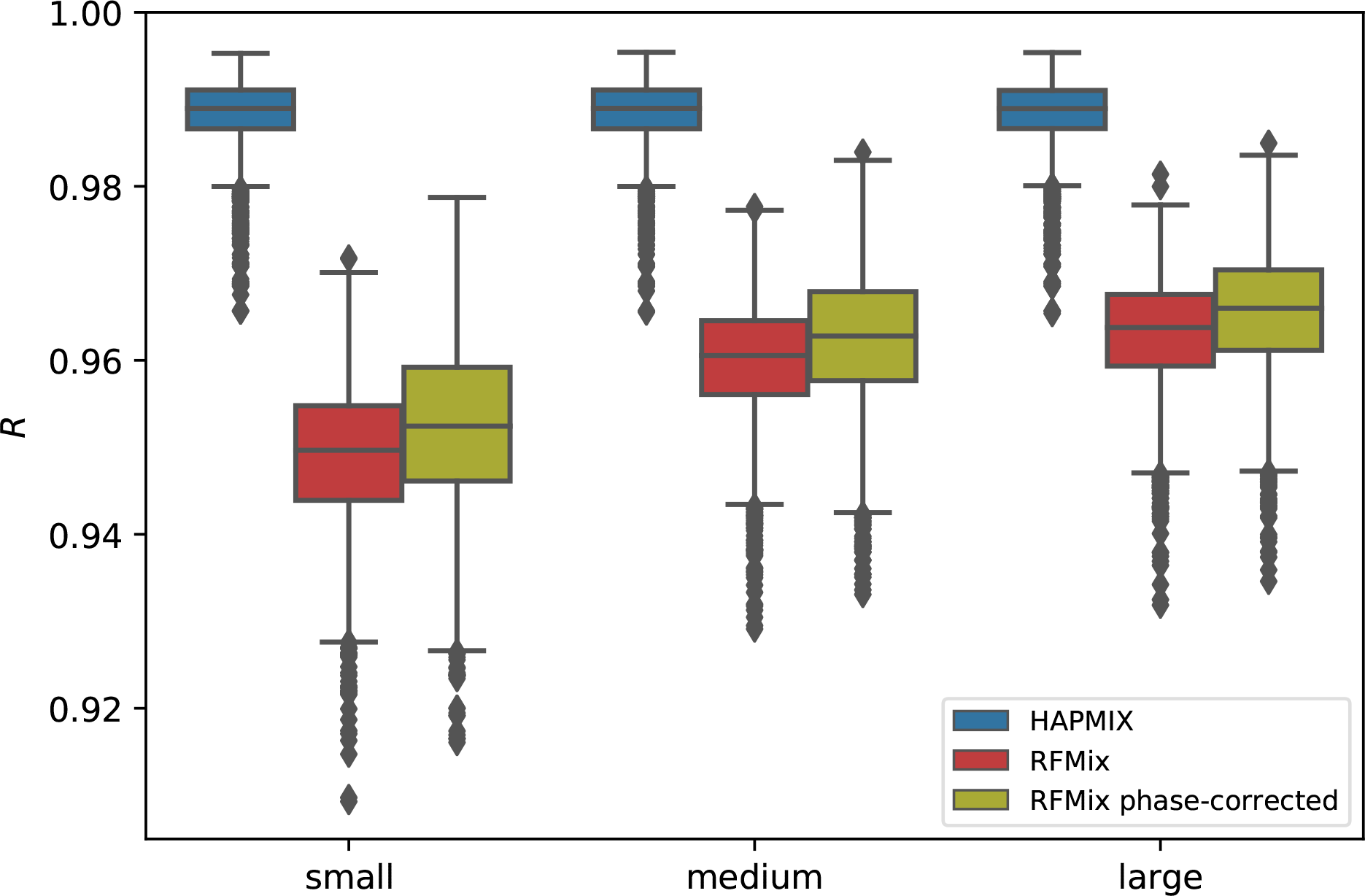
Phase correction with TRACTOR yields only a small improvement in phase-based local ancestry inference. Plot shows correlations between inferred and true local ancestry assignments *R* by phasing sample size. The RFMix phase-corrected method uses the output from TRACTOR. The plot includes data points from all other simulation variables.

### Admixture time estimates using phase-based LAI are strongly biased

To characterize the impact of LAI accuracy on downstream applications, we calculated admixture time estimates using the output of both FLARE and HAPMIX. Because HAPMIX produces unphased local ancestry calls, we provided these as input to PAPI ^20^ and report its time estimates (see Methods). Figure 5 plots the deviations of the admixture time estimates from the ground truth (*T*_*dev*_ = *T*_*inferred*_ − *T*_*true*_) for the reference phasing strategy data and for each phasing sample size. Strikingly, estimates using HAPMIX’s calls are virtually unbiased, while the FLARE-based estimates are strongly upwardly biased, even when the phase quality is high. Specifically, the average values of *T*_*dev*_ using FLARE are 14.2, 11.1 and 9.84 for small, medium and large sample sizes, respectively. HAPMIX showed a slight downward bias of -0.0909 for all sample sizes (recall that HAPMIX is a phase-free LAI method and its results are expected to remain unchanged with respect to phasing sample size).

**Figure 5:**
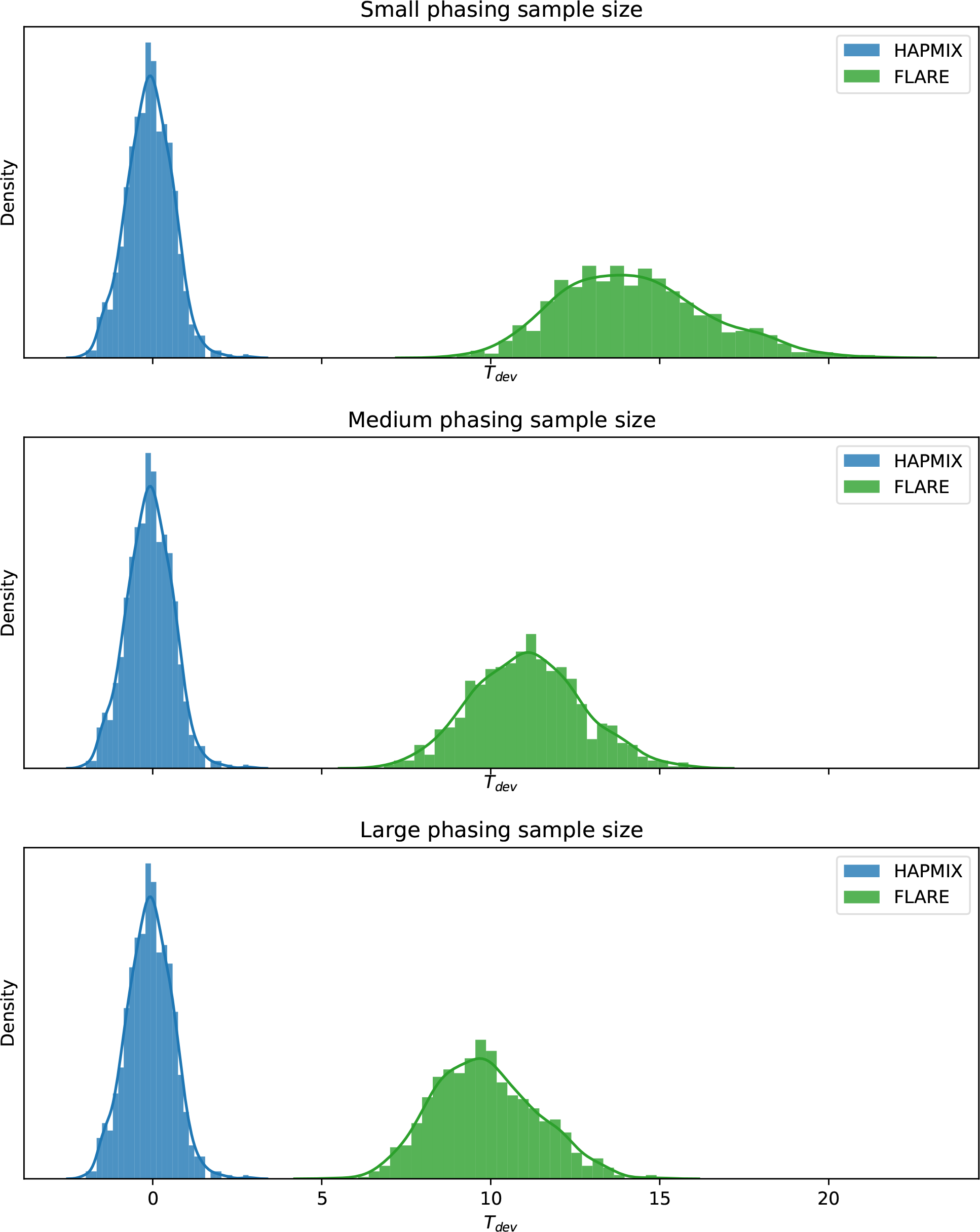
Deviations of admixture time estimates using local ancestry calls from HAPMIX and FLARE. Plots depict histograms of the deviations of admixture time estimates for individual samples for (top) small, (middle) medium, and (bottom) large phasing sample sizes. Data points are pooled across all other simulation variables.

## Discussion

Here we analyzed the performance of four state-of-the-art LAI methods and found that despite large sample sizes and good quality phase, phase-based methods do not perform as well as phase-free ones in terms of call accuracy of unphased ancestry states (Figure 3). Furthermore, unphased LAI remains more effective even after applying TRACTOR’s phase-correction algorithm, which identifies likely switch errors from changes in the local ancestry on both haplotypes of an individual (Figure 4).

Based on our analyses, the most important factor that affects phase quality in admixed individuals is sample size (Figure 2). Moreover, for a fixed sample size, we found empirically that it is best to include the unadmixed reference haplotypes when phasing, with a slight improvement by conditioning on them (using, e.g., SHAPEIT’s `--reference` or Beagle’s `ref` option ^27^) rather than by pooling them with the admixed individuals (as in the rephasing strategy). That is, there is an improvement in phase quality alone by including these reference haplotypes—perhaps because they increase the sample size. Additionally, conditioning on reference haplotypes or rephasing them makes the target individuals’ phase more consistent with these panels and is therefore recommended for LAI and other downstream processes (such as imputation) that utilize reference panels.

A combination of both switch errors breaking up ancestry tracts and reduced accuracy of phase-based LAI methods can substantially impact downstream analyses that leverage these data, such as estimating admixture times. Our findings show that even with good phase quality (i.e., phasing sample sizes of 5810 individuals), FLARE’s admixture time estimates are biased by more than an order of magnitude relative to those based on HAPMIX calls (Figure 5). Indeed, concern over phase-based LAI quality motivated us to develop PAPI for estimating time since admixture and parent ancestry proportions using unphased local ancestry calls ^20^. Because of these issues, most estimates of time since admixture necessarily use trio-phased data, but given the quality of phase-free methods, tools such as PAPI can leverage the more abundant non-trio samples to perform even sample-specific admixture time estimates.

The difference in accuracy between phase-based and phase-free LAI is substantial enough that careful deliberation on whether to apply phase-based LAI may be warranted, depending on the setting. Our results suggest that HAPMIX in particular is considerably more accurate than the other methods in all parameter settings (see Results), making it a clear choice when high quality LAI calls are required but high quality phase is unavailable. Of course, a key limitation of HAPMIX is that it applies only to two-way admixed samples; yet LAMP-LD is a competitive alternative (*R* = 0.978 on average versus HAPMIX’s *R* = 0.988) that can perform LAI in multi-way admixed samples.

Notably the sample sizes we used yield good but far less accurate phase than is available with biobank-scale data ^24,28^ and phase-based LAI may perform competitive with—or better than—phase-free methods given such well-phased data. (Indeed, a previous benchmarking study that used the phase produced by their simulation pipeline found that RFMix outperformed LAMP-LD ^18^, while in our statistically phased samples, LAMP-LD performs better.) Further, we focused on two-way admixed samples, examining the archetypal case of admixture between relatively divergent populations (African and European) where we can expect the highest quality LAI. In that regard, our work can be viewed as a best-case empirical analysis for sub-biobank-scale data, and demonstrates that challenges for phase-based methods apply even in such settings.

This work highlights important concerns that can arise in LAI from phase quality issues—even when phasing is done with several thousand genotyped samples. Future LAI methods development could focus on new phase-free approaches or on modeling and correcting switch errors within phase-based methods; the most relevant switch errors for LAI accuracy likely change the local ancestry of the underlying haplotype, and efforts to correct these could have a huge impact on the accuracy of resulting local ancestry calls. Overall, our findings draw attention to the importance of designing studies that apply LAI carefully, with due consideration to factors such as phase quality, utilization of reference panels, choice of LAI method, and admixture demography. Most crucially, we demonstrate that the phase of the admixed samples has a large impact on LAI accuracy for phase-based methods, an issue implicitly mitigated by phase-free LAI methods.

## Acknowledgements

Funding for this work was provided by NIH grant R35 GM133805. Computing was performed on a cluster administered by the Biotechnology Resource Center at Cornell University. Samples and data of The Charles Bronfman Institute for Personalized Medicine (IPM) BioMe BioBank used in this study were provided by The Charles Bronfman Institute for Personalized Medicine at the Icahn School of Medicine at Mount Sinai (New York). Phenotype data collection was supported by The Andrea and Charles Bronfman Philanthropies. Funding support for genotyping, which was performed at The Center for Inherited Disease Research (CIDR), was provided by the NIH (U01HG007417). The datasets used for the analyses described in this manuscript were obtained from dbGaP at http://www.ncbi.nlm.nih.gov/gap through dbGaP accession number phs000925.v1.p1.

