## Supplemental Figures 1 for "Phase-free local ancestry inference mitigates the impact of switch errors on phase-based methods"

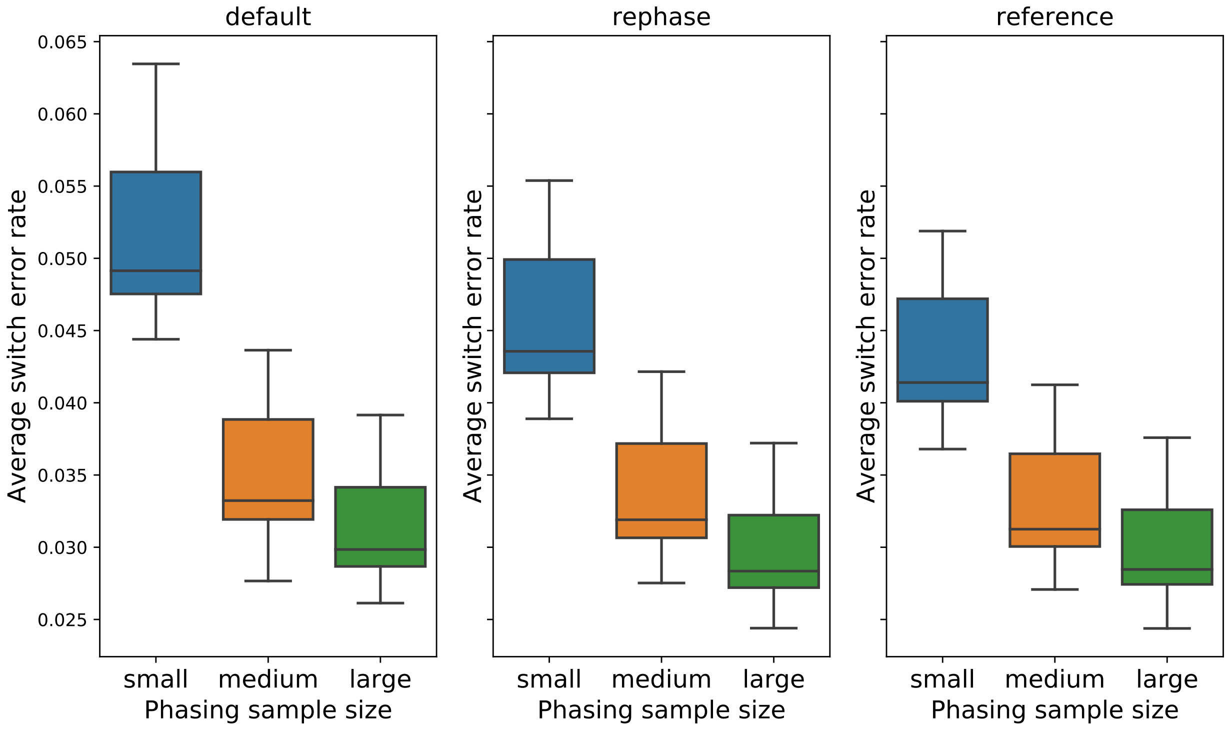


Figure S1: **Switch error rates in simulated samples for different panel phasing strategies.** Each panel shows switch error rates for each phasing sample size for one of the panel phasing strategies (default, rephase, or reference). Each panel includes data points from all other simulation variables (i.e., African ancestry proportion and time since admixture).

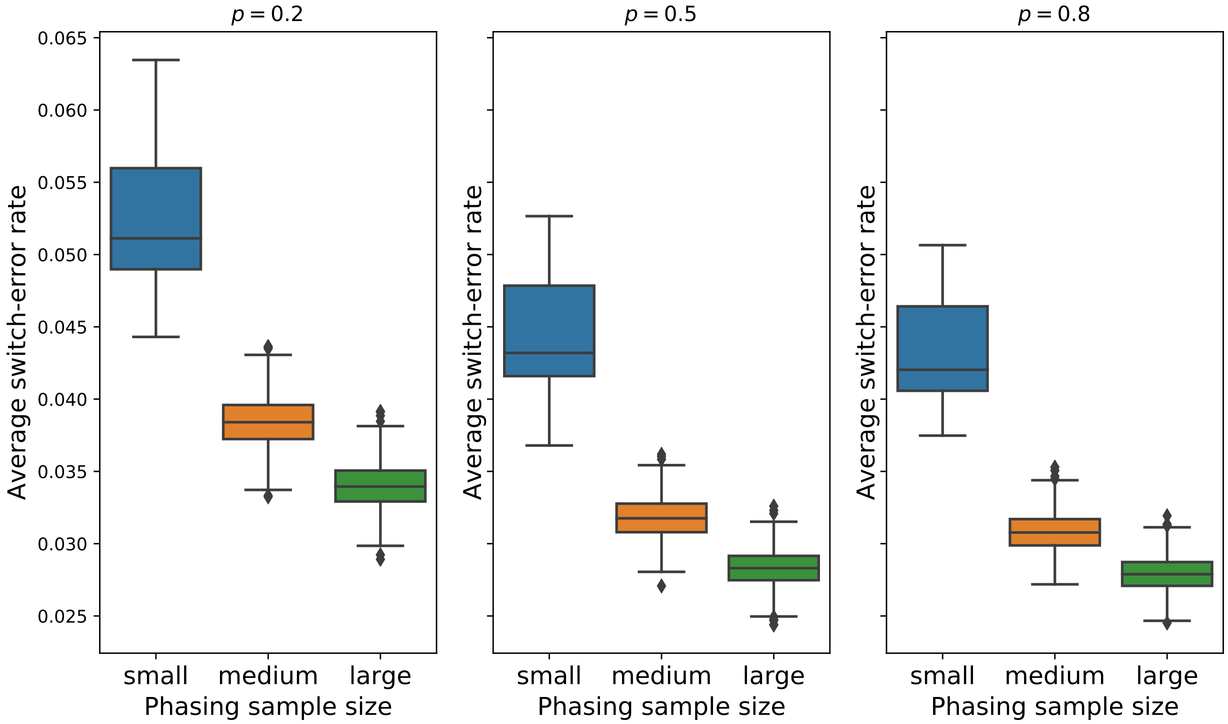


Figure S2: **Switch error rates in simulated samples for different African ancestry proportions.** Each panel shows switch error rates for each phasing sample size for one of the African ancestry proportions ($p = 0.2$, $p = 0.5$, or $p = 0.8$). Each panel includes data points from all other simulation variables (i.e., panel phasing strategy and time since admixture).

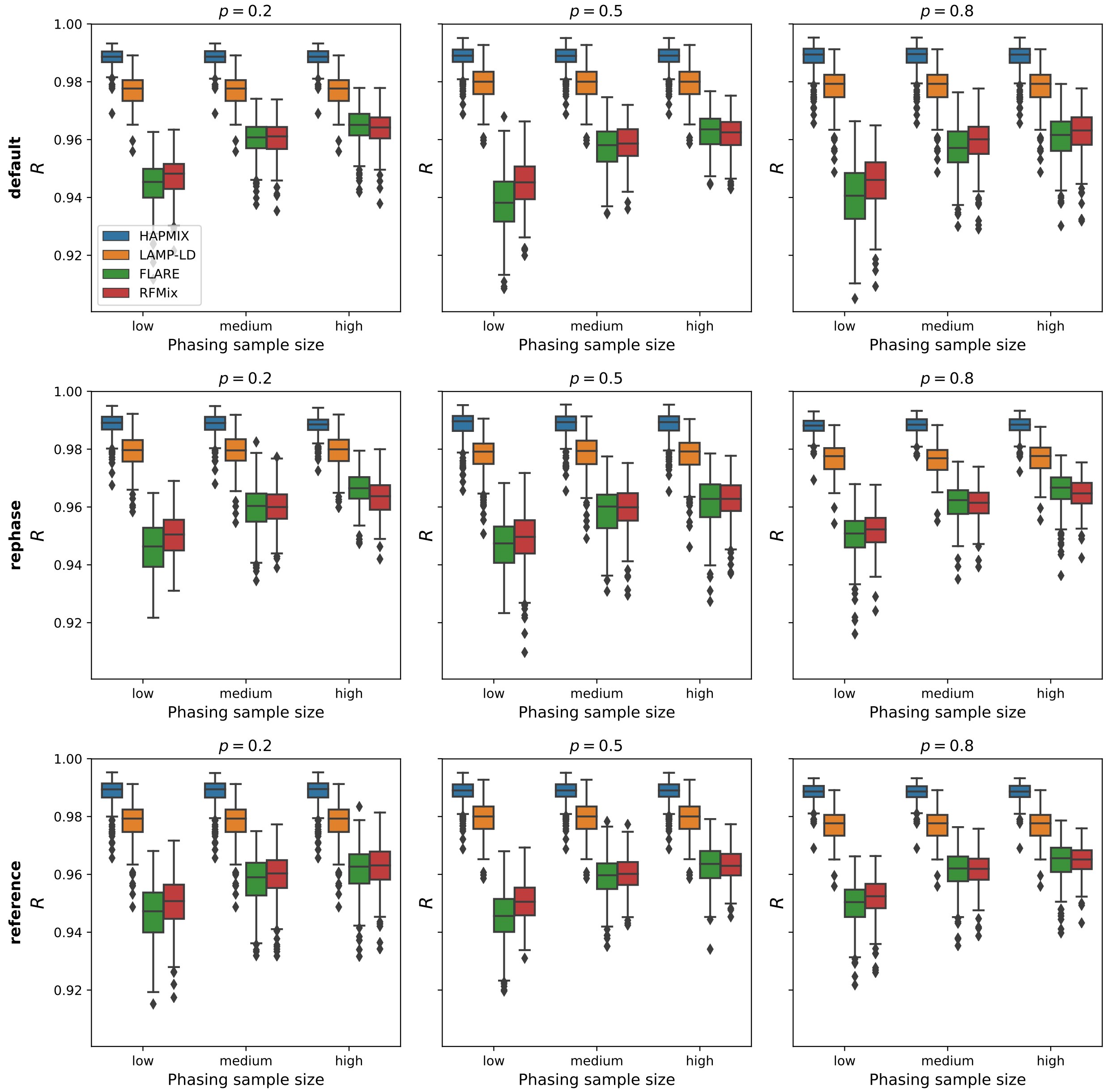


Figure S3: **Performance of local ancestry inference methods across different panel phasing strategies and African ancestry proportions.** Each panels shows boxplots of correlations between inferred and true local ancestry assignments $R$ for each phasing sample size arranged in a grid of panel phasing strategy (default, rephase, or reference) by African ancestry proportion ($p = 0.2$, $p = 0.5$, or $p = 0.8$). Each panel includes data points from the other simulation variable (time since admixture).

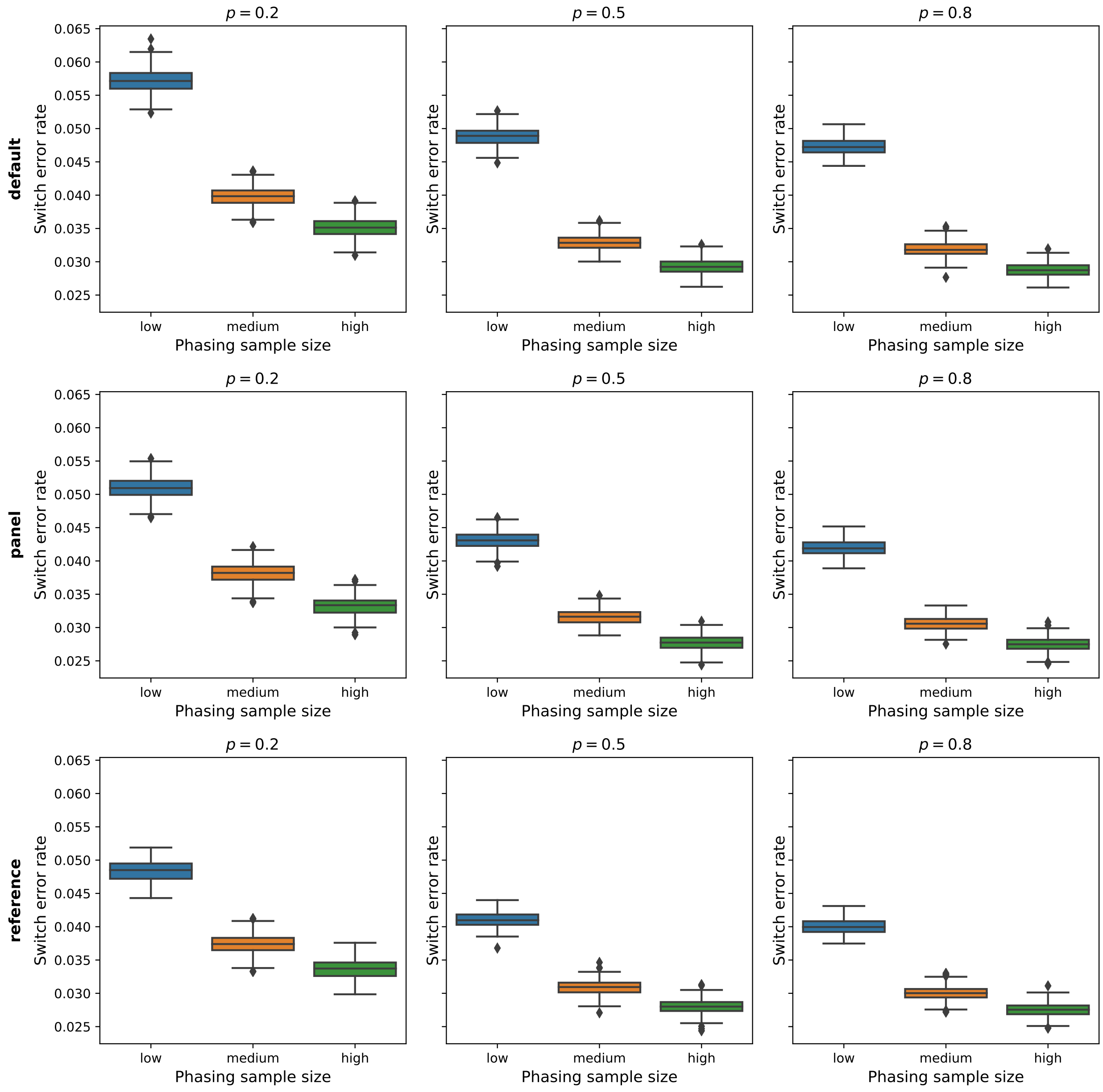


Figure S4: **Switch error rates across different panel phasing strategies and African ancestry proportions.** Each panel shows switch error rates for each phasing sample size arranged in a grid of panel phasing strategy (default, rephase, or reference) by African ancestry proportion ($p = 0.2$, $p = 0.5$, or $p = 0.8$). Each panel includes data points from the other simulation variable (time since admixture).

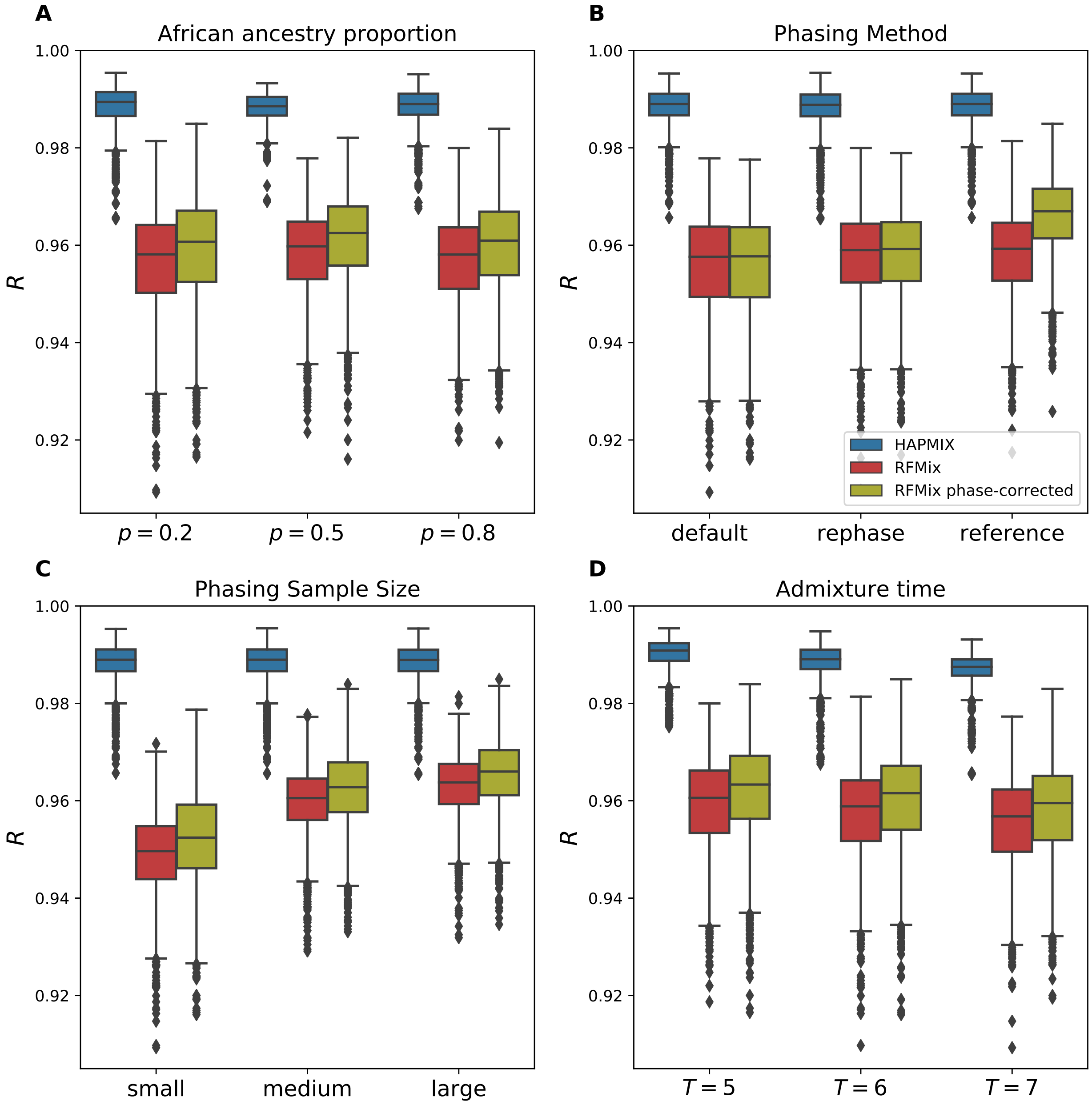


Figure S5: **Performance of HAPMIX, RFMix and phase-corrected RFMix across all simulation parameters.** Correlations between inferred and true local ancestry assignments $R$ plotted against (A) proportion of African ancestry $p$, (B) panel phasing strategy (C) phasing sample size, and (D) admixture time $T$ (i.e., generations since admixture). Each panel includes data points from all other simulation variables.
